# Deciphering the astrocytic contribution to learning and relearning

**DOI:** 10.1101/2024.01.12.572819

**Authors:** Lorenzo Squadrani, Carlos Wert-Carvajal, Daniel Müller-Komorowska, Kirsten Bohmbach, Christian Henneberger, Pietro Verzelli, Tatjana Tchumatchenko

## Abstract

Astrocytes play a key role in the regulation of synaptic strength and are thought to orchestrate synaptic plasticity and memory. Yet, how specifically astrocytes and their neuroactive transmitters control learning and memory is currently an open question. Recent experiments have uncovered an astrocyte-mediated feedback loop in CA1 pyramidal neurons which is started by the release of endocannabinoids by active neurons and closed by astrocytic regulation of the D-serine levels at the dendrites. D-serine is a co-agonist for the NMDA receptor regulating the strength and direction of synaptic plasticity. Activity-dependent D-serine release mediated by astrocytes is therefore a candidate for mediating between long-term synaptic depression (LTD) and potentiation (LTP) during learning. Here, we show that the mathematical description of this astrocytic regulation is consistent with the classic Bienenstock Cooper Munro (BCM) model for synaptic plasticity, which postulated the existence of an activity-dependent LTP/LTD threshold. We show how the resulting mathematical framework can explain the experimentally observed behavioral effects of astrocytic cannabinoid receptor knock-out on mice during a place avoidance task and give rise to new testable predictions about the learning process advancing our understanding of the functional role of neuron-glia interaction in learning.

## Introduction

Our understanding of synaptic plasticity, a fundamental brain process underlying learning and memory, has greatly advanced through a synergistic combination of experimental research and theoretical modeling. Among the notable theories in this field, the Bienenstock, Cooper, and Munro (BCM) theory stands out [1, 2]. Built on a concise set of assumptions, this theory accurately describes the development of pattern selectivity in cortical neurons [3, 4]. Furthermore, it has successfully predicted phenomena that were yet to be observed at the time of its conception, guiding experiments to uncover them [5–7]. However, how this form of plasticity could be implemented in the brain is not fully understood yet [2, 8].

The core postulate of the BCM theory is the existence of a dynamic threshold for long-term potentiation (LTP) which slides as a super-linear function of post-synaptic activity. Several hypotheses have been proposed in the last decades to give a physiological justification for this postulate [7, 9–12]. Many of them involve intracellular calcium dynamics in the post-synaptic neuron. However, how intracellular calcium and excitability of neurons are dynamically regulated to enable LTP and LTD in the parameter domains observed in experiments remains a matter of debate. At the same time, a growing body of experiments points at astrocytes and extracellular D-serine regulation as previously overlooked candidates for closing the gap between abstract plasticity models, neurobiology, and behavioral output.

Astrocytes interact with neurons via a complex system of neuroactive substances, and they have been implicated in various neuronal functions, including synaptic transmission, synaptic plasticity, and memory formation [13–19]. D-serine is thought to be the main endogenous ligand for the co-agonist binding site of N-methyl-D-aspartate receptors (NMDARs) in the hippocampus [20, 21], and occupation of this site is required to enable long-term plasticity [22, 23]. It has been observed that extracellular D-serine levels are regulated by astrocytes and determine NMDAR tone and NMDA-dependent plasticity [14, 24–26]. In some experiments, D-serine was shown to affect the activity-dependence of long-term synaptic changes, in a manner similar to the shifting of a potentiation threshold, resembling that of the BCM theory [27]. More recently, it was discovered a new mechanism for the astrocytic regulation of synaptic available D-serine [28], characterized by a bell-shaped dependence on the frequency of the post-synaptic activity. The mechanism exhibits remarkable similarities to the core postulates of BCM theory and is shown to impact behavioral choices in mice when it is disrupted. Could this newfound astrocytic mechanism be the missing link that unites the BCM theory with the intricate biology of the brain? In this paper, we explore the hypothesis that the astrocytic regulation of D-serine represents the physiological implementation of BCM plasticity in the brain; we refer to this as the “D-serine hypothesis”. To support it, we present a formal derivation of the BCM rule, starting from the mathematical description of the astrocytic mechanism observed by Bohmbach et al. [28]. Building upon the hypothesis, we develop a mathematical model capable of explaining precise behavioral effects resulting from astrocytic manipulation, as observed in experiments, and produce new testable predictions on the learning process. Overall, our results point to new, compelling ideas to reduce the gap between our theoretical understanding of plasticity and our experimental knowledge of neuron-glia interactions in learning.

## Results

### The D-serine hypothesis: astrocytes orchestrate BCM plasticity

Recent experiments on mice CA1 hippocampal neurons have shed light on a novel molecular mechanism, forming a feedback loop through which the activity of pyramidal neurons influences their future dynamics and plasticity [28]. The feedback loop is mediated by astrocytes and is enacted by two main molecules: endocannabinoids released by neurons upon activation, which bind to the cannabinoid receptors (CBRs) of astrocytes [29, 30], and D-serine, released into the extracellular environment in response to astrocytic calcium signaling triggered by CBRs activation [14, 24, 31, 32] (Fig. 1a).

**Fig. 1.**
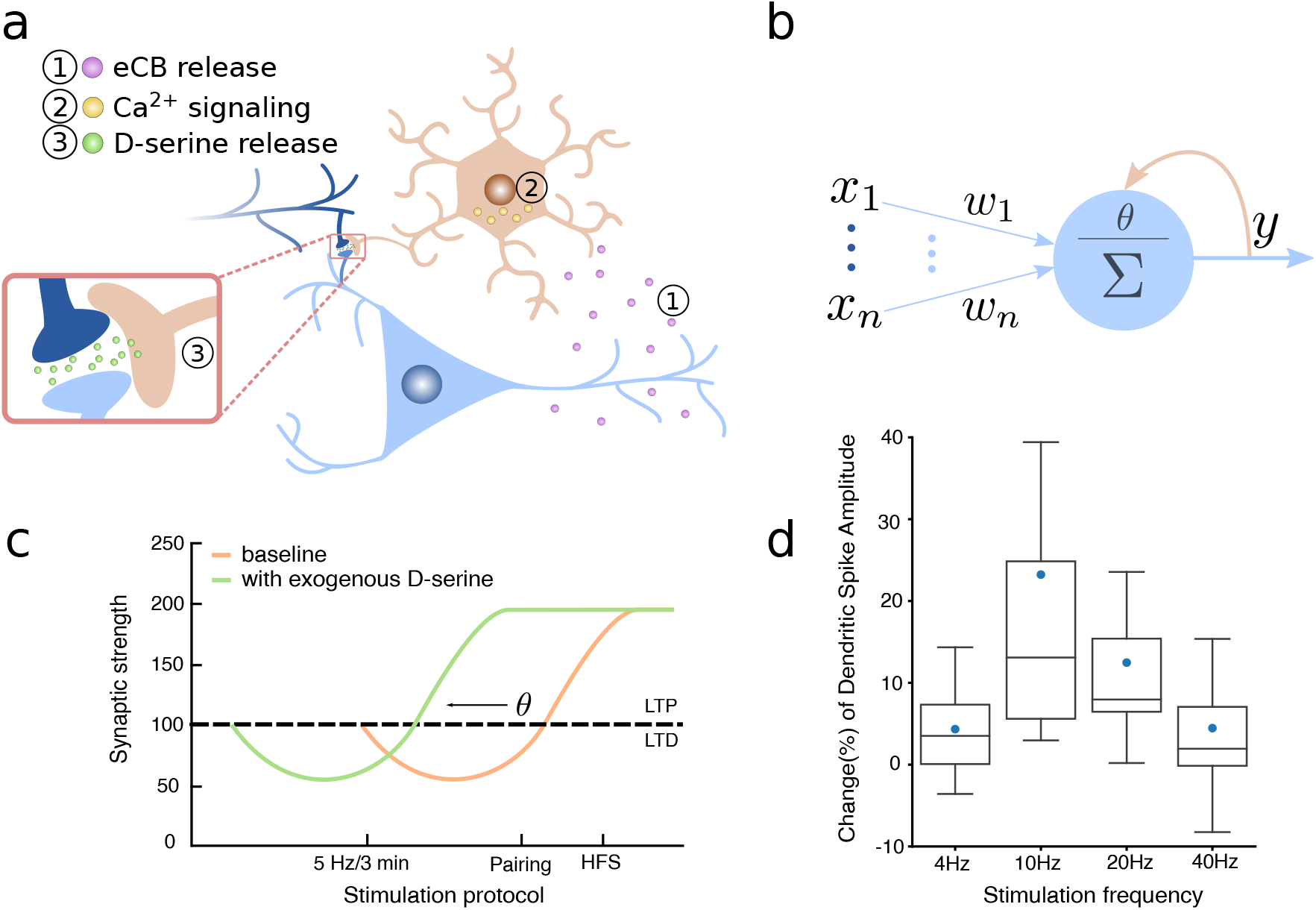
The D-serine hypothesis. (a,b) The astrocytes-mediated feedback loop observed by Bohmbach et al. [28] shows remarkable similarities with the BCM postulate of a synaptic modification threshold *θ* sliding as a function of the post-synaptic activity *y*. The feedback loop is described in three steps: (1) endocannabinoids are released by the post-synaptic neuron upon activation, (2) they bind with astrocytic cannabinoid receptors and trigger an intracellular calcium signaling, which results in (3) the release of D-serine at the neuron dendrites. (c) The increase (decrease) of D-serine concentration at the synapse promotes LTP (LTD), which can be interpreted as the shifting of a threshold for LTP, like the BCM one (figure adapted from Panatier et al. [27]). (d) The astrocytic regulation of D-serine presents a peculiar bell-shaped dependence on the frequency of post-synaptic stimulation, analogous to the super-linear dependence of the BCM’s threshold on the post-synaptic activity. Changes in the D-serine level are indirectly measured through changes in the amplitude of dendritic spikes’ slow component (figure adapted from Bohmbach et al. [28]).

Let *y* represent the average firing rate of a neuron population, and *d* denote the concentration of D-serine in the extracellular environment. Based on previous observations, we describe the dynamics of *d* so that it follows a given function of the post-synaptic activity, denoted as *D*(*y*), with a time constant *τ*_*d*_:

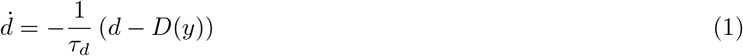

The right-hand side of Eq. (1) can be interpreted as the sum of a D-serine degradation/uptake term and a term that describes activity-dependent D-serine supply. The function *D*(*y*) describes how post-synaptic activity influences D-serine levels. The dependence of the D-serine release on the post-synaptic activity has been studied by monitoring D-serine-dependent dendritic integration during the axonal stimulation of CA1 pyramidal neurons at different frequencies [28]. Notably, an unexpected bell-shaped dependence has been observed, with a peak at 10*Hz* (Fig. 1, d). Based on this, we choose the simple mathematical form:

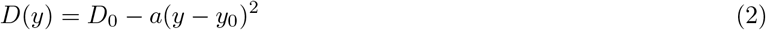

where *D*_0_ is the maximum level of D-serine, occurring when *y* = *y*_0_, and *a* is a constant with the appropriate dimensions.

The importance of D-serine stems from its interaction with synaptic NMDAR, which plays a fundamental role in the induction of synaptic plasticity [33]. D-serine has been identified as an essential endogenous ligand for the glycine site of synaptic NMDARs in the brain [20, 21, 27]. Consequently, its presence is required for the activation of NMDARs. Consistent with these findings, alterations in D-serine levels have been unequivocally associated with changes in the activity-dependent nature of long-term synaptic modifications [27]. In particular, the same stimulation protocol induced LTP or long-term depression (LTD) depending on the level of D-serine. In the presence of a high (low) level of D-serine, LTP (LTD) is facilitated, and the same effect is equally seen with three different stimulation protocols. Such observation suggests that the effect of D-serine alterations can be interpreted as a shifting of a threshold for LTP induction (Fig. 1c). We denote with **w** a vector of synaptic strengths and with **x** a vector of pre-synaptic activities. We define the integration of the input vector with the synaptic strengths to track the neural activity with respect to the reference value *y*_0_, thus *z* = *y* − *y*_0_ = **w** · **x**. Finally, we define an update rule for the synaptic strengths in such a way that the direction of plasticity depends on the level of D-serine through a function *θ*(*d*):

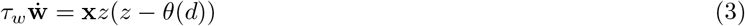

where *τ*_*w*_ is a time constant. The proportionality of the update to the pre- and post-synaptic firing realizes the Hebbian principle, while the quadratic dependence on the post-synaptic rate has been suggested by experimental measurements ([5, 27]) and has been derived from spikes-based learning rules (STDP [34, 35], calcium-based plasticity [36]). The equation is formally equivalent to the BCM update rule [1], with the important difference that here the threshold is no more a postulated dynamical variable but a function of the D-serine concentration. We show now that, choosing the function *θ*(*d*) according to experimental observations and considering the dynamics of D-serine described by Eq.s 1 and 2, we obtain a dynamical system equivalent to BCM.

According to the fact that a higher level of D-serine facilitates LTP, we choose the threshold function to be inversely proportional to the D-serine concentration:

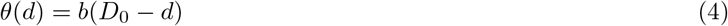

where *b* is a constant with the appropriate dimensions. The threshold is 0 when *d* is at its maximum value *D*_0_, and it increases as *d* decreases. We define the new variable 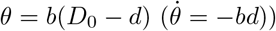, and substitute it in the Eq.s 1 and 3, obtaining:

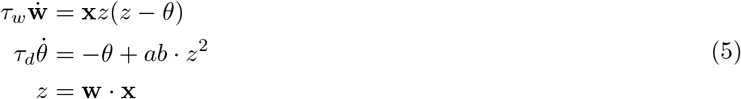

which is equivalent to the BCM plasticity rule [1], and coincides exactly by setting *ab* = 1, as we do for the rest of this work.

### Testing the D-serine hypothesis within a behavioral context

The biological interpretation of BCM proposed above allows us to explore how the disruption of the D-serine feedback loop changes learning performance. In mice, this intervention has been shown to affect learning performance during a place avoidance task [28]. The original experiment (Fig. 2) consisted of a mouse freely exploring an O-shaped maze, triggering an air puff each time it passed through a fixed position; after the mouse learned to avoid that position, the puff was moved to the opposite side. Learning was assessed by measuring the number of visits to each position. The disruption of the D-serine feedback loop corresponded to a deletion of astrocytic type 1 CBRs in a group of mice (mutant mice). In the initial phase of the task, mutant mice learned to avoid the air puff correctly, en par with the control mice (Fig. 2b). However, when the puff position was switched, the mutant showed *slower learning* with respect to control mice. The mutation thus has a significant and specific impact on behavior, which cannot be described as a generic learning impairment, offering an ideal test bench for our hypothesis.

**Fig. 2.**
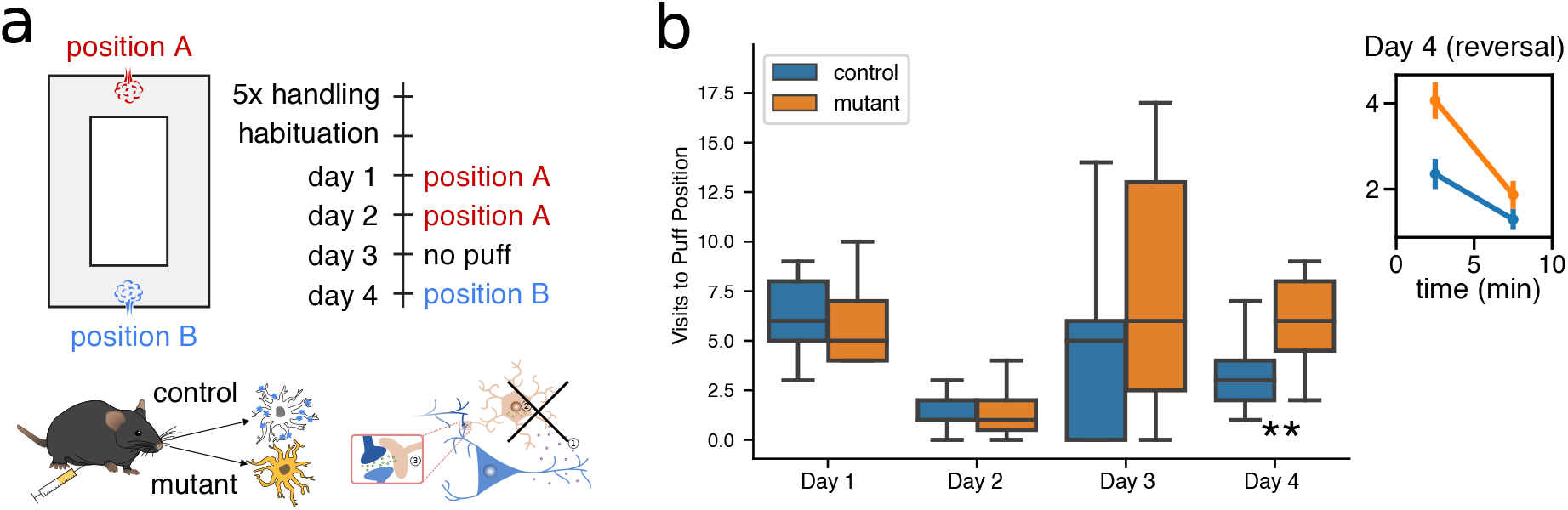
Learning impairment of mutant mice during a place-avoidance taks. (a) The learning ability of the mouse was experimentally tested with a place-avoidance task driven by an aversive stimulus. The mouse was free to explore an O-shaped maze, triggering an air puff each time it passed through a fixed location; after the mouse learned to avoid that position, the puff was moved to another location. The test is repeated with two groups of mice: a control group, and a group in which the astrocytic type 1 cannabinoid receptors had been knocked out (mutant). (b) Mutant mice showed slower learning with respect to control mice, only during reverse learning (Day 4). Figures adapted from [28].

To approach this problem within the theoretical learning framework described in section 5, we model a simplified version of the task with a two-state environment, *S*^1^ and *S*^2^ (Fig. 3). Such a description is suitable because the only relevant measurements in the original experiment were the number of visits to the initial and the final positions of the puff. For a detailed description of how the agent moves between the states refer to the section *Methods*. The simulation is divided into two phases. At the beginning, a punishing signal is associated with *S*^2^ (Phase 1); the agent has to learn to avoid the punishing state. After a given amount of time has elapsed, the punishing signal is moved to *S*^1^ (Phase 2), so that the agent needs to learn the new configuration, reverting the previously learned behavior. Learning emerges from the adaptation of the synaptic weights of a single action unit, which is implemented with the BCM plasticity model, modified in order to account for the punishing signal.

**Fig. 3.**
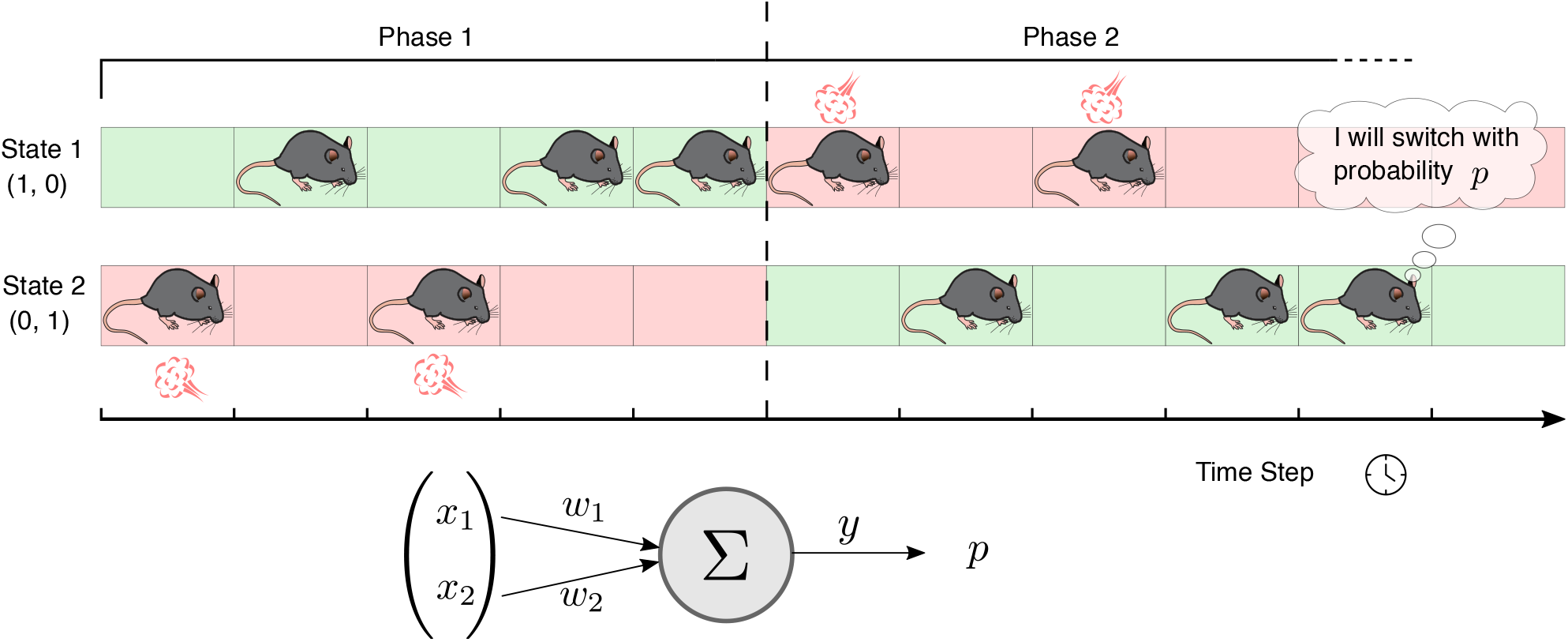
Simulation of the passive place-avoidance task. In our simulation of the experiment, we simplify the movement of the mouse as a transition between two locations (State 1 and State 2 in the figure), one of which is associated with a punishment (red background). The simulation is divided into two phases. During the first phase State 2 is punishing, while in the second phase the punishment is moved to State 1. At each time step, the mouse decides whether to stay in the current state or to switch. The decision is based on the current state, represented by a one-hot encoding state vector, and on the information the mouse has about the states, encoded in the weights *w*_1_ and *w*_2_ of a single action unit. The integration of the state vector with the synaptic weights determines the activity *y* of the unit, which in turn determines the probability *p* of switching state.

### Extending BCM to solve a reinforcement-driven task

The original BCM model, as presented by Bienenstock, Cooper, and Munro [1], is a purely unsupervised learning rule. Consequently, it is insufficient for behavioral learning tasks that require distinguishing between neural activities associated with positive or negative outcomes. It is widely acknowledged that such distinction is mediated in the brain through the actions of neuromodulators, such as dopamine [37, 38]. In the realm of theoretical modeling, neuromodulation has been described by extending two-factor learning rules (where the factors are pre- and post-synaptic activities) to *three-factor* learning rules, with the third factor being a reinforcement-signaling variable [39]. While this problem has been addressed for general Hebbian learning and Spike Timing Dependent Plasticity [40–43], it has not been explored for BCM. Here, we propose a simple and natural model that addresses this issue, allowing for the solution of the punishment-driven place avoidance task described in the previous section.

To this aim, we first rewrite the system 5 for a discrete time step, in consistency with the literature on reinforcement learning [44]; on each transition from state *S*_*t*_ to state *S*_*t*+1_, the unit having an activity *y*_*t*_ determined by the weights **w**_*t*_ and the input **x**_*t*_ = **x**(*S*_*t*_), the BCM update rule reads:

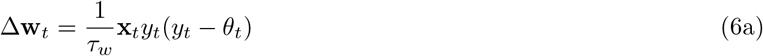

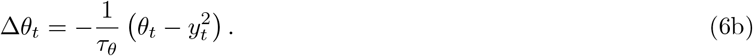

Following the ideas discussed above, we introduce a third, reinforcement factor in Eq. 6a:

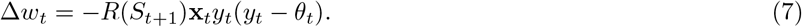

Note that the value of the reinforcement factor −*R*(*S*_*t*+1_) depends on the destination state, providing information about the outcome of the action which was taken at time *t*. The weight update thus depends on quantities distant in time. This is a common problem in reinforcement learning (distal reward problem [41]), and there has been considerable effort to understand how such time gap could be bridged in the brain. The most common solution is postulating the existence of an “elegibility trace” at the synapses, tracing previous activity and leading to a weight change only in the presence of a third, reinforcement factor [39]. Experimental evidence in support of eligibility traces on the time scale of seconds has been collected in the last years (for a review, see [39]).

We refer to the resulting plasticity rule as ‘Reinforcement-BCM’ (R-BCM). From a functional perspective, BCM plasticity allows the action unit to develop selectivity with respect to the incoming stimuli, thus making the agent select one of the two states. The extension to the R-BCM allows the agent to select one specific state according to the values of the reinforcement factor, and to change selection when the values are switched. Thus, the new rule can solve the behavioral task described in the previous section. Further details and the analysis of stability are provided in *Methods*.

### Simulations reproduce learning deficit in mutant mice

The learning deficit observed by Bohmbach et al. [28] following the disturbance of neuron-astrocyte communication exhibits distinct characteristics, which present challenges in directly mapping it to astrocytic and neuronal activity. Naively incorporating the effect of the disruption of D-serine regulation as a generic learning impairment (for example by linking it to a larger time scale of the weight dynamics) would not align with the experimental findings, as mutant mice exhibited a deficit only in the second phase of learning, when the puff position is switched. Importantly, our biological interpretation of BCM allows us to give a more precise description of the disruption of the D-serine feedback loop, by modifying the dynamics of D-serine. Experiments show that, after the deletion of astrocytic type 1 CBRs, the D-serine levels are not affected anymore by neuronal stimulation. The concentration of D-serine is thus constant (non-zero because this would completely prevent NMDA-dependent plasticity). This aspect is described by Eq.2 by setting a zero value of the activity 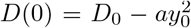. In practice, this corresponds to a constant value for the LTP threshold *θ* = *θ*_0_.

The mice moving in the two-state environment and utilizing R-BCM for synaptic weight updates learn to avoid the state associated with the negative reinforcement (Fig. 4, top). In the initial phase of the simulation, states *S*^1^ and *S*^2^ are linked to positive and negative reinforcement signals, respectively. Consequently, the frequency of state visits *S*^1^ increases, occurring approximately 9 out of 10 times. The same behavioral pattern is observed when the LTP threshold is fixed, simulating the disruption of the D-serine feedback loop in mutant mice. This behavior aligns with the experimental findings in Bohmbach et al. [28].

**Fig. 4.**
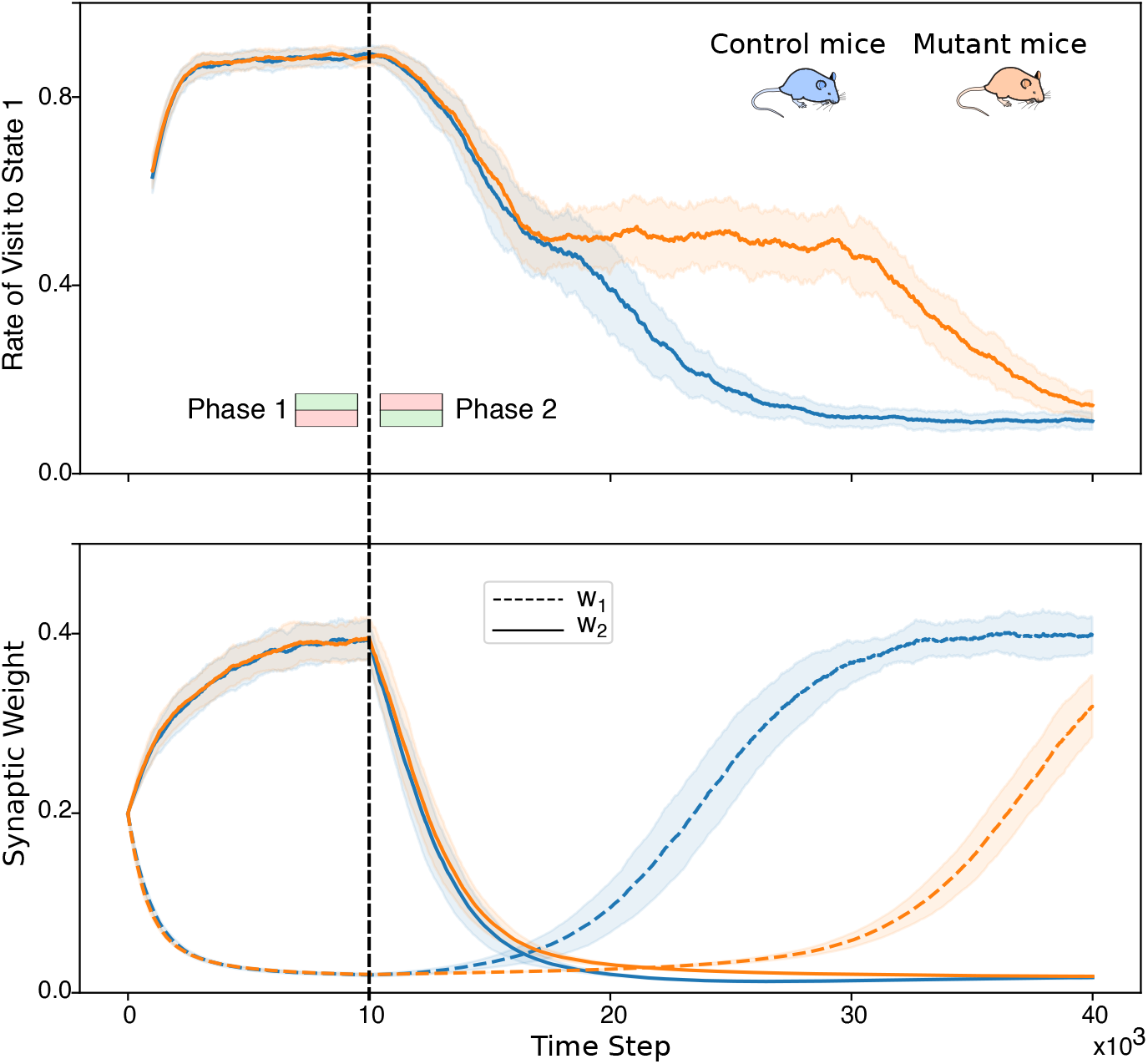
The model reproduces learning deficit in mutant mice. The figure shows the results of the simulated place-avoidance experiment (described in Fig.3) for control mice and mutant mice. The top graph shows the rate of visit to the state *S*^1^, computed as the relative number of visits to state *S*^1^ in the last 1000 time steps. The bottom graph shows the evolution of the synaptic weights. All the curves are averages over 50 simulated mice. During the first phase of the simulation (left of the black line) the states *S*^1^ and *S*^2^ are associated with a positive and negative reinforcement signal, respectively. Coherently, the frequency of visits of the mouse to state *S*^1^ increases to about 9 times out of 10. Both weights converge to a finite value, one of them close to zero, the other close to 0.4. There is no difference in learning between control mice and mutant mice. During the second phase (right of the black line), the reinforcement signals are switched. Both groups of mice learn to avoid the state *S*^1^, with a visit frequency of about 1 time out of 10. However, mutant mice show a significant delay during this phase of reverse learning. During this phase, the weights switch their values. The switching is faster in control mice (top) than in mutant mice (bottom). More precisely, the higher weight decays with the same speed in both groups, reflecting the initial drop in the visiting frequency (top figure, second phase). However, the low weight increases earlier and faster in control mice than in mutant mice.

The frequency that each state is visited is determined by the synaptic weights of the action unit, as described in the section *Methods*. In the bottom graph of Fig. 4, we plot the weights’ evolution throughout learning. During Phase 1, the weight *w*_1_, which represents the probability of switching state from *S*^1^ to *S*^2^ (the punishing state), correctly converges to a value close to zero. This means that the probability of *going to* the punishing state is low. The weight *w*_2_, which represents the probability of switching state from *S*^2^ to *S*^1^, converges to a higher value, but lower than 0.5. This means that the probability of *going away from* the punishing state is not close to 1, as might be intuitively expected from a probabilistic agent that learned to avoid this situation. However, this property can be justified and interpreted as proof of the coherence of the model. The *avoidance* behavior can be conceptually divided into two sub-behaviors: *going away from* the aversive situation and *not going back to* it. Such a distinction is important because the two behaviors are unlikely to arise from the same neurobiological processes. For instance, the *not going back to* behavior requires memory of past experiences, while the *going away from* does not since the aversive stimulus is present and can drive the behavior directly. Coherently, because we are only modeling one mechanism (synaptic plasticity), the model does not capture the combination of both behavioral components. Irrespective of these considerations, the experimentally determined behavior and the learning differences are reproduced correctly.

After switching the reinforcement signals associated with the states (second phase of the simulation), the control mice quickly adapt their behavior to avoid the other state. In contrast, the mutant mice learn significantly slower (Fig. 4, top). Importantly, such difference in the two phases of learning is again in agreement with what has been observed in [28] (Fig. 2).

An inspection of the evolution of the weights reveals interesting insights into the deficit (Fig. 4, bottom). The weight associated with the previously punishing state drops down without significant difference in both classes of mice. However, the weight associated with the newly punishing state rises much slower in the mutant mice. The reason for that lies in the blockade of the D-serine regular dynamics, as seen by looking at the D-serine levels during reverse learning (Fig. 5, left). In control mice, reverse learning is accelerated by a temporary increase in the D-serine levels, which results in a lower LTP threshold.

**Fig. 5.**
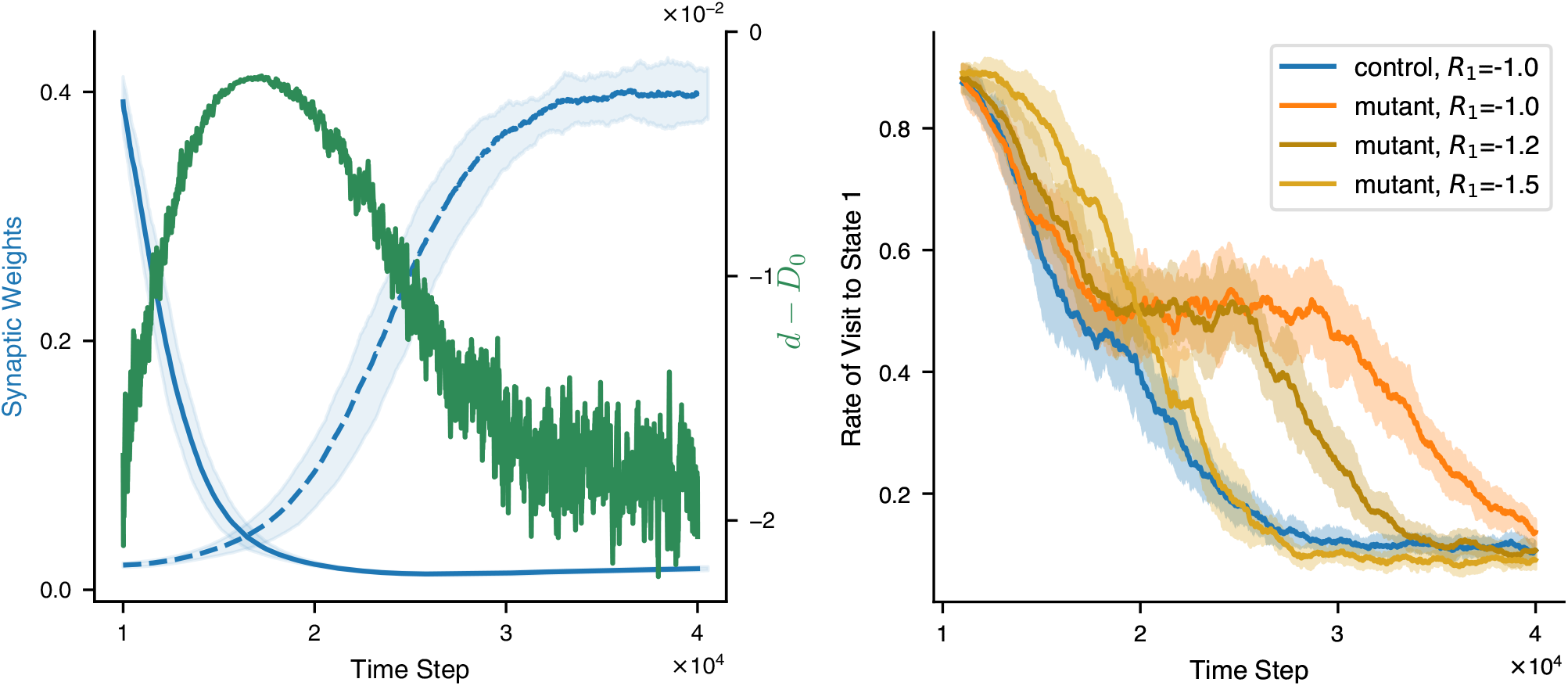
(left) Increased D-serine levels during reverse learning. The blue lines are the evolution of the two synaptic weights during reverse learning of normal mice (same lines of Fig. 4, bottom graph, second phase). The D-serine concentration relative to its maximum value *D*_0_ is superposed (green line). The scale for *d* − *D*_0_ is of course different from that of the synaptic weights, and it is drawn on the right axis. The D-serine levels are increases during reverse learning and they return to the basal value when the switching is completed. **(right) Increasing the aversiveness of the stimulus recovers learning speed for mutant mice**. The rate of visit to state *S*^1^ during the second phase of learning is plotted, for control mice with *R*_1_ = *−*1, and for mutant mice with *R*_1_ = *−*1, *−*1.2, *−*1.5. The mutant mice learn faster as the magnitude of the *R*_1_ increases, and it reaches the speed of the control mice for *R*_1_ = 1.5.

Next, we tested whether increasing the intensity of the punishing signal (by modulating the values *R*_*i*_, see the section *Methods*) for the mutant mice could restore the learning speed. In Fig. 5, right side, we plot the rate of the visit to state *S*^1^ during the second phase of learning for control mice with *R*_1_ = −1, and for mutant mice with *R*_1_ = −1, −1.2, −1.5. The mutant mice learn faster as the magnitude of the *R*_1_ increases, and it reaches the speed of the healthy mice for *R*_1_ = −1.5.

## Discussion

Here, we have proposed a new hypothesis for the physiological realization of BCM-like plasticity in the brain. The hypothesis is based on an astrocyte-mediated feedback loop in CA1 pyramidal neurons, initiated by the release of endocannabinoids by post-synaptic neurons and closed by D-serine level regulation at the dendrites. This mechanism was only recently discovered by Bohmbach et al. [28], and it permitted us to elaborate a formal derivation of BCM based on brain biology. The involvement of D-serine in synaptic plasticity and its possible relation with the BCM rule was already suggested in experimental literature [27, 45], but it has not found appropriate consideration in theoretical modeling yet. Of particular relevance is the fact that the existence and the dynamics of the modification threshold have been derived from experimental data rather than postulated. Notably, the new formulation includes and describes the role of astrocytes and gliotransmitter in synaptic plasticity and has therefore important consequences in connecting the learning model to behavior and to the role of astrocytes in learning. Indeed, it allowed us to simulate the disruption of the astrocytic feedback loop, which has been observed to impact the behavior of mice learning to avoid an aversive stimulus in a dynamic environment. Simulations within this theoretical framework produced a behavior strikingly similar to the observed one. Our results strongly suggest that modeling activity-dependent D-serine dynamics is a crucial step towards more biological plausible versions of the BCM rule, and plasticity models in general.

The most significant advantage of establishing a direct connection between a mathematical model of learning and the biological structures and molecules in the brain is the ability to generate new testable predictions. The learning curves depicted in Fig. 5 predict that an increase in the extracellular D-serine level should be observed during the reverse learning phase. When confirmed by experiments, this result can point at an interesting functional role for the D-serine regulatory system, i.e. ensuring a fast plasticity response to changes in the external environment. The evolution of the synaptic weights during reversal learning in the mutant mice (Fig. 4, bottom) suggests that the learning impairment is due to the inhibition of LTP caused by astrocytic CBR knockout. The inhibition is not homogeneous across the range of synaptic strength however: in the first phase of learning, when the synaptic strength starts from an intermediate value, control mice and mutant mice do not show any difference in the LTP; in the second phase, when synaptic strength starts from a highly depressed value, LTP is slower in mutant mice compared with control mice. The BCM model originated as a theory of synaptic plasticity in visual cortex. Here, we proposed a biological implementation of BCM based on experimental manipulations of astrocytes and D-serine dynamics in the CA1 region of the hippocampus. Therefore, observing similar properties in the visual cortex would be another interesting confirmation of the D-serine hypothesis. Experiments that deny or confirm such properties are missing in the current literature, as to our knowledge.

The D-serine hypothesis for the adaption of the BCM threshold considers the extracellular dynamics at the synapse, while the previously studied hypotheses are concerned with the intracellular dynamics, usually of calcium [10], but also of other molecules like the kinase CamKII [12, 46]. For this reason, our hypothesis does not directly contradict the previous ones. On the contrary, the extracellular D-serine dynamics might very well represent the cause of alteration of the intracellular dynamics which ultimately lead to the regulation of the potentiation threshold. The interaction between the extracellular synaptic dynamics and the intracellular dynamics, crucially mediated by the NMDARs dynamics, deserves further investigation and could lead to a coherent and more general model of synaptic plasticity, extending the mathematical framework proposed by Shouval et al. [10].

The BCM plasticity theory has been related to Spike-Timing Dependent Plasticity (STDP), by showing that a BCM-like rate-based plasticity can arise from STDP through an averaging operation [34]. In this context, the threshold for synaptic potentiation depends on the shape of the STDP window. According to the D-serine hypothesis, the threshold is function of the synaptic D-serine concentration. To understand the relation and the compatibility of the two models it is thus crucial to understand how different D-serine levels affect the STDP window. Such a question could be tackled experimentally, but also theoretically by modeling how NMDARs shape the STDP window. Gjorgjieva et al. a model of STDP where the synaptic change window depends on the time of the last post-synaptic spike preceding the plasticity-inducing spikes pair (triplet STDP); interestingly, they showed that there is a tight correspondence between this triplet STDP rule and the BCM rule.

The standard Hebbian learning rule is known to be susceptible to stability issues [48]. Intense neural activity leads to synaptic potentiation, which in turn amplifies future activity. This process is sometimes referred to as the Hebbian positive feedback mechanism. The BCM theory addresses the problem by introducing the sliding threshold mechanism [6, 11], which can be implemented in the brain through the D-serine feedback loop. However, it has been observed that learning can still occur even when the D-serine feedback loop is disrupted, suggesting the existence of alternative forms of stabilization. The Reinforcement-BCM model proposed in this work successfully reproduces the disruption of the D-serine feedback loop and offers an alternative source of stability through the zero-averaging of positive and negative reinforcement signals. However, this solution has some inherent limitations. It is not easily applicable in the absence of reinforcement or when only positive or negative reinforcement is present. Additionally, it provides stability on average, which restricts the learning speed as excessively rapid weight updates could compromise the system’s averaging operation and lead to instability.

When the BCM mechanism of stabilization based on a sliding potentiation threshold was initially proposed, experimental research was conducted to investigate the existence of such a biological mechanism [6, 49]. The studies aimed to determine whether the recent history of synaptic activity could influence the induction of LTP. It was observed that the induction of LTP under a specific stimulation protocol was not fixed but could indeed be strongly influenced by the prior synaptic activity. However, the influence on LTP induction was found to be specific to the stimulated synapses rather than generalized to all inputs onto the postsynaptic neuron, as postulated by the BCM theory. A similar contradiction arises from our reformulation of the BCM model. We assumed that the D-serine level, represented by the variable *d* in Eq. 1, is the same at all synapses, which led to a neuron-wide potentiation threshold identical to BCM. However, considering the close proximity of astrocytes to dendrites, it is possible that their regulatory action on D-serine can target individual or specific groups of synapses, posing a contradiction to the neuron-wide potentiation threshold assumption in the BCM model. This possibility will be further explored when extending the D-serine hypothesis to a network of neurons, possibly providing new insights on the intricated interactions between neurons and glial cells.

## Methods

### Model of the place-avoidance task

We simulate a mouse performing a place avoidance test with an aversive stimulus. The spatial environment is modeled as a discrete two-state space *S*^1^, *S*^2^. At each time step, the agent chooses between two actions: staying in the current state or moving to the other one. The states are represented by one-hot encoding vectors **x**^1^ = (1, 0) and **x**^2^ = (0, 1). The vectors are used as synaptic inputs for a single action unit, describing the decision process of the mouse. If at time step *t* the mouse occupies the state *S*_*t*_ with associated input vector **x**_*t*_, the activation of the action unit is given by the scalar product of the input vector with its synaptic weights:

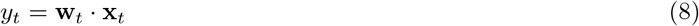

The initial values of the weights are (*w*_1_, *w*_2_) = (0.2, 0.2) and the agent start from a random state. The mouse switches state at the next time step with a probability *p*_change_ directly proportional to the firing rate of the action unit, with a minimum value of 0.05:

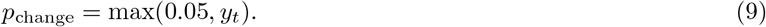

The lower bound of *p*_*change*_ was introduced to maintain a minimum level of exploration. After each transition, the synaptic weights and the synaptic threshold (i.e. the D-serine level) are updated:

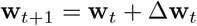

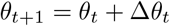

The learning rule consists of the BCM plasticity rule modified in order to account for the negative reinforcement (see section 5 and the rest of Methods).

### Learning rule

On each transition from state *S*_*t*_ to state *S*_*t*+1_, the unit having an activity *y*_*t*_ determined by the weights **w**_*t*_ and the input **x**_*t*_ = **x**(*S*_*t*_), the weights are updated according to the following learning rule:

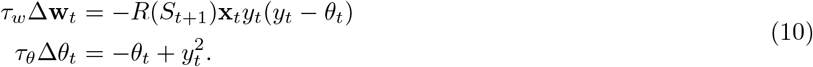

This is the BCM learning rule with the addition of multiplicative factor depending on the final state. The equation for the weight change is dissected into four possible cases (the four possible transitions) in Table 1. We refer to the rule as “R-BCM”.

**Table 1.**
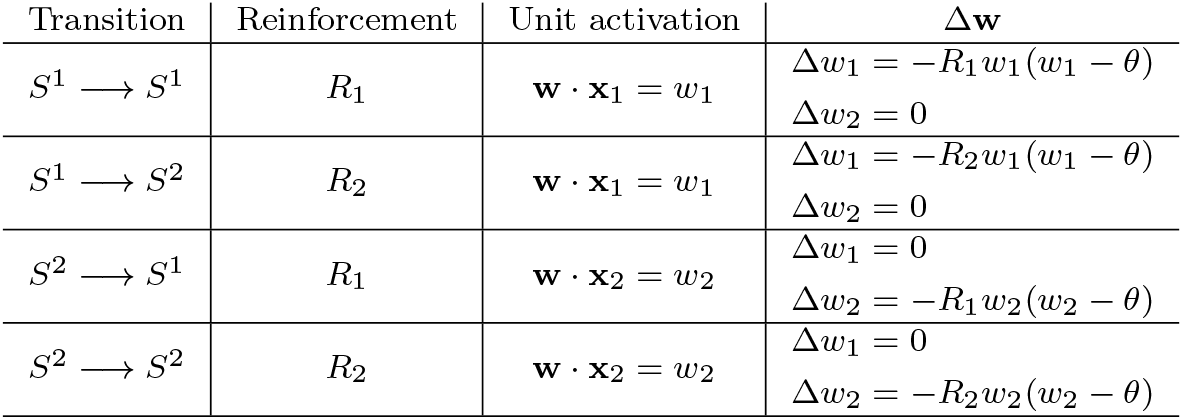
R-BCM dynamics. At each step, the agent performs one of the state transitions in the first column of the table. The reinforcement signal is determined by the final state. The action unit activity is determined by the initial state. The synaptic weights are updated according to Eq.7. We note that since Δ**w** is proportional to the one-hot input vectors, only one of the weights changes at each transition.

We compute the stationary points of R-BCM with the same procedure used for obtaining the stationary points of the original BCM [8, 50], i.e. replacing the stochastic differential equation (7) with the same equation averaged over the input environment:

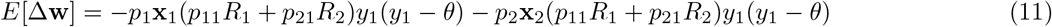

where *y*_1_ = **w** · **x**_1_, *y*_2_ = **w** · **x**_2_, *p*_*i*_ is the probability of being in state *i*, and *p*_*ij*_ is the probability of the transition from state *j* to state *i*. The dynamics of the agent can be described as a Markov chain over the two-state space, so that the probabilities **p** = (*p*_1_, *p*_2_) obey **p**(*t* + 1) = Π**p**(*t*), where Π is the transition matrix. We recall that the probability of switching state is related to the firing rate of the action unit through Eq. 9. Thus, for a fixed weight vector, the transition matrix reads:

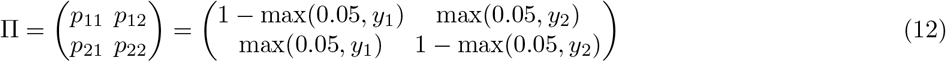

The transition matrix is thus a function of the weight vector **w** (through *y*_*i*_ = **w** · **x**_*i*_). However, since we are looking for the stationary weight vectors, we assume the transition matrix to be constant. In this case, the probability vector **p** will converge to the eigenvector of Π with eigenvalue 1, which is:

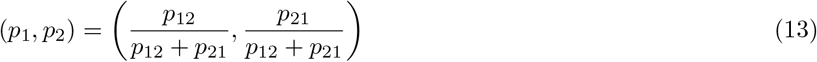

The stationary points are computed by solving the equation *E*[Δ**w**] = **0**. Assuming that the input vectors **x**_1_ and **x**_2_ are linearly independent (as it was in our simulations), this leads back to solving two equations independently:

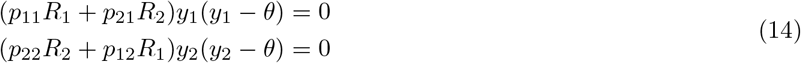

We obtain 7 distinct stationary points:

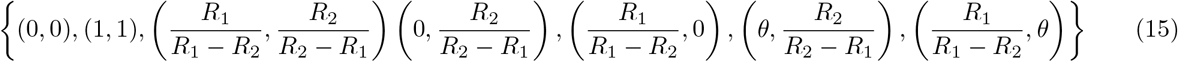

where the value of *θ* is the determined by the equation (valid in the “slow-learning” approximation [50], i.e. *τ*_*w*_ ≫ *τ*_*θ*_):

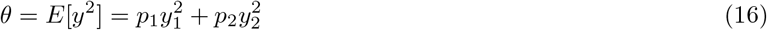

where *p*_1_ and *p*_2_ are given by Eq.13. We note that the coincidence of the stationary points of the deterministic equation 11 with the stationary points of the stochastic equation 6a has to be proven, by showing that the solutions converge. This was done analytically for the BCM equation [51], but doing the same for the R-BCM is non trivial. Instead, we check the coincidence of the stationary points through numerical simulations. We use numerical simulations also to check the stability properties of the points, and find that only one is stable, according to what we expected by looking at the average vector field (Fig. 6). The stable point is:

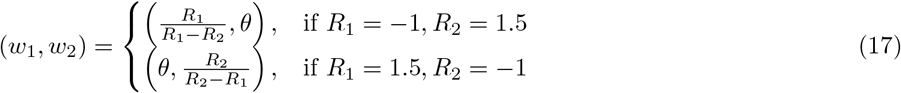

**Fig. 6.**
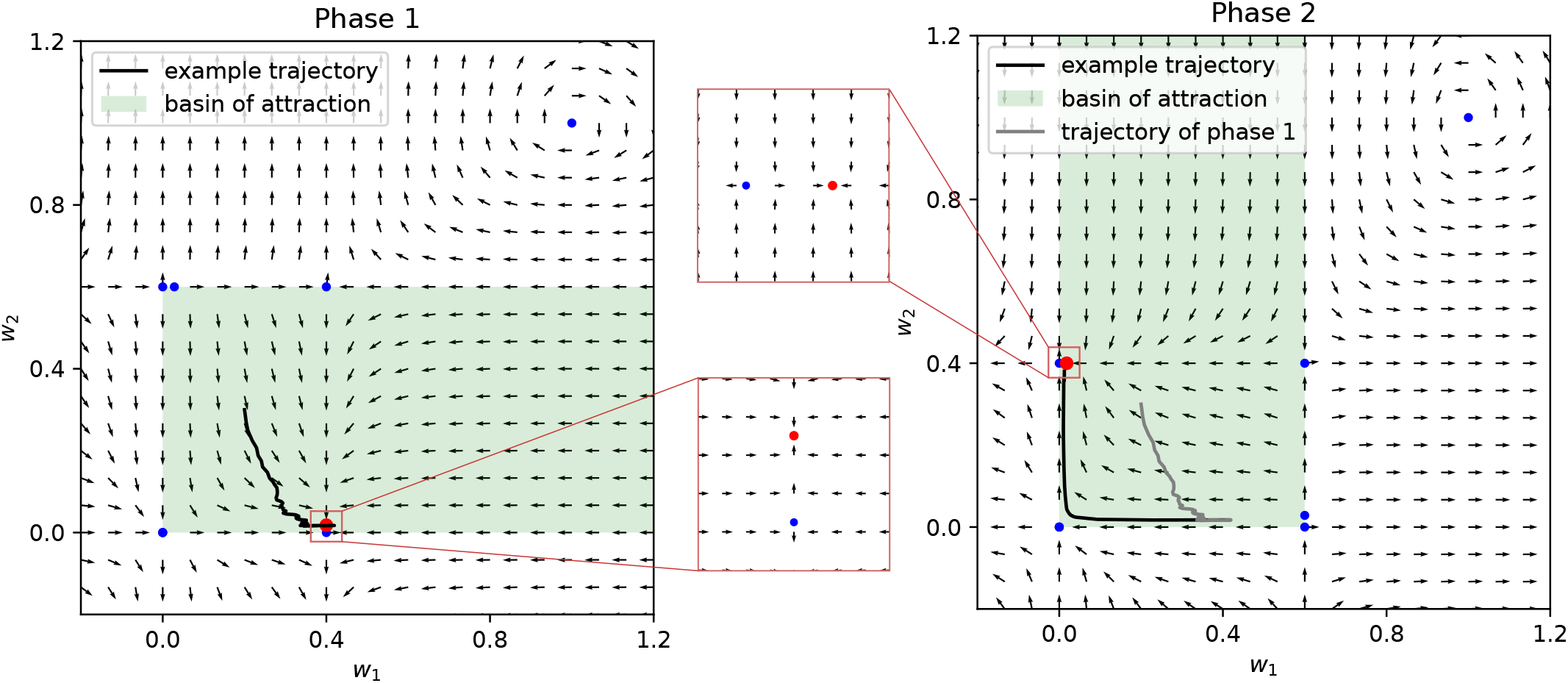
Stability of the R-BCM system. The stationary points (Eq.15) and the average vector field (Eq.11) are plotted in the weight space, during the first phase of learning (left, *R*_1_ = −1 and *R*_2_ = 1.5) and during the second phase (right, the values of *R*_*i*_ are switched). The blue dots denote unstable stationary points, the red dots denote stable stationary points. The average vector field *E*[Δ**w**] reveals that there is only one stable point (insets). The green area denotes the basin of attraction of the stable point. The stability properties are checked also through simulations with multiple initial conditions. An example trajectory is plotted, showing that the weights tend towards the stable point during phase 1, and then correctly tend towards the new stable point after the switching. We note that the stationary points and the average vector field of phase 1 (before switching) and phase 2 (after switching) are related by a reflection across the identity line. It is important that the initial stable point falls inside the basin of attraction of the new stable point after the reflection. This is ensured by the fact that the positive *R*_*i*_ is greater in magnitude than the negative one, as can be seen by substituting the values in Eq.15.

Importantly, as in the original BCM, this is a *selective* point, meaning that the activity *y*_*i*_ = **w** · **x**_*i*_ is high for one input and low for the other.

### Parameters

For the time constants of the system (10), we set *τ*_*w*_ = 100 and *τ*_*θ*_ = 50, with intrinsic time units. Numerical simulations show that it must be *τ*_*w*_ ≫ 1 in order for the stationary points of Eq. 17 to be stable. This is due to the fact that the value of *R* in Eq. 10 changes at every time step Δ*t* = 1, and the stability of the system relies on an averaging operation over the *R* (see Eq. 14). Concerning *τ*_*θ*_, the derivation of the stationary points in the previous section is valid under the assumption *τ*_*w*_ ≫ *τ*_*θ*_, so that Eq. 16 holds. However numerical simulations show that a stable point exists also for greater values of *τ*_*θ*_. Indeed, a stable point exists also when *τ*_*θ*_ → +∞, which correspond to the case of the mutant mice. In this case, however, the stable point is not given anymore by Eq.16. Numerical simulations show that the point is very close to the original one.

The function *R* can assume only two values, *R*_1_ = *R*(*S*^1^) and *R*_2_ = *R*(*S*^2^). During the first phase of training, we set *R*_1_ = 1.5 and *R*_2_ = −1; the values are switched during the second phase. With this values, the stable steady state is such that the agent choices tend to avoid the state *S*_*i*_ with *R*_*i*_ = −1. One might thus be tempted to interpret −1 as a negative reinforcement and 1.5 as a positive or neutral reinforcement. However, it is important to be aware that such interpretation is arbitrary: the avoidant behavior is determined by the *combination* of the two values. The fact that the values are opposite in sign ensures that the stationary points in (15) lay in the positive quadrant. The fact that the positive value is greater in magnitude than the negative value, ensures that, immediately after the switching, the system is in the basin of attraction of the new stable point (see Fig. 6).

When simulating the disruption of the D-serine feedback loop in mutant mice, the LTP threshold is fixed at *θ*_0_ = 0.02. This value correspond to the average stationary value of the threshold obtained with normal mice, far away from the switching time.

## Code Availability

All the code used to simulate the experiments will be made publicly available upon publication. In the review stage, an implementation of the model and visualisation of the experimental outcomes in Netlogo[52] can be found at: https://verzep.github.io/R-BCM.html

## Author Contributions

All the authors conceived the study and initiated the work. L.S., P.V, T.T. formalized the mathematics of the model and designed the learning task. L.S. simulationed the model and analyzed the data. All authors contributed to the manuscript and took part in the scientific discussion of the results.

## Acknowledgement

This research was supported by University of Bonn Medical Center, the Deutsche Forschungsgemeinschaft (DFG, German Research Foundation) – Project-ID 227953431 – SFB 1089 (PV, CWC, DMK, TT, CH), and the iBehave Network, which is funded by the Ministry of Culture and Science of the State of North Rhine-Westphalia (LS, CH,TT).

